# Co-Registration of Optoacoustic Tomography and Magnetic Resonance Imaging Data from Murine Tumour Models

**DOI:** 10.1101/636035

**Authors:** Marcel Gehrung, Michal Tomaszewski, Dominick McIntyre, Jonathan Disselhorst, Sarah Bohndiek

**Affiliations:** Department of Physics, University of Cambridge, UK; Cancer Research UK Cambridge Institute, University of Cambridge, UK; Werner Siemens Imaging Center, Preclinical Imaging and Radiopharmacy, University of Tuebingen, Germany

**Keywords:** Optoacoustic Tomography, Magnetic Resonance Imaging, Image Registration

## Abstract

As optoacoustic tomography emerges as a mainstream preclinical imaging modality, understanding the relationship between optoacoustic and other imaging biomarkers in the context of the underlying tissue biology becomes vitally important. For example, assessment of blood haemoglobin concentration and oxygenation can be achieved using OT, and also by several magnetic resonance imaging (MRI)-based techniques. To evaluate the relationship between these metrics and the relative performance of the two modalities in assessment of haemoglobin physiology, co-registration of their output imaging data is required. Unfortunately, this poses a significant challenge due to differences in the data acquisition geometries. Here, we present an integrated framework for registration of OT and MR image data in small animals. Our framework combines a novel MR animal holder, to improve animal positioning for deformable tissues, and a landmark-based software co-registration algorithm. We demonstrate that our protocol significantly improves registration of both body and tumour contours between these modalities.

## 1. Introduction

Optoacoustic tomography (OT) is an emerging imaging modality, able to reveal the distribution of tissue optical absorption coefficient in real-time with a spatial resolution of ∼ 180 µm at ∼ 3 cm penetration depth [1]. Thanks to the distinct optical absorption profiles of oxy- and deoxyhaemoglobin, acquiring OT data at multiple wavelengths (multispectral optoacoustic tomography, MSOT) makes it possible to derive optoacoustic imaging biomarkers that relate to total haemoglobin concentration (THb) and blood oxygenation (SO_2_) [2]. Application of these functional MSOT imaging biomarkers has been shown to provide insight into both the architecture and function of the blood vasculature, for example, in cancer imaging, where it can be used to monitor tumour development [3, 4] and detect response to therapy [5, 6].

Functional imaging of the blood vasculature is also possible with a wide range of magnetic resonance imaging (MRI)-based techniques. Taking the example of cancer imaging, dynamic contrast enhanced (DCE) MRI [7], blood oxygen level dependent (BOLD) [8], oxygen enhanced (OE) MRI [9] and arterial spin labelling (ASL) MRI [10] have all been demonstrated to provide insight into tumour blood vessel function and the surrounding tissue hypoxia. The question thus arises, how do these different imaging techniques compare with OT and do their imaging biomarkers correlate?

The correct combination of spatial information from different imaging modalities requires careful alignment of the images and hence an efficient co-registration algorithm. This is usually achieved in both patient and small animal imaging by careful body positioning and scanning process optimisation, aided by software-based alignment. Well-established, clinically used solutions are available [11, 12] and provide excellent results for fusion of positron emission tomography (PET), computed tomography (CT) and MRI data. Unfortunately, modalities such as OT that involve different scanning geometries and positioning of the animal or patient pose a significant challenge to co-register. Successful co-registration of OT and MR images has been reported previously in the brain of small animals [13, 14], however, being contained within the skull, the brain is not subject to any deformation due to external forces, making it a relatively simple organ to co-register.

Here, we present a new integrated framework for registration of MSOT and MR image data in pre-clinical studies of small animals, which can be applied to soft, deformable tissues such as tumours. The method combines a novel animal holder design and a robust co-registration algorithm. We first describe the method and show its performance for co-localization of the internal tumour structure between the modalities. We then demonstrate the improvement in co-registration achieved by the combination of hardware and software-based solutions, compared to the manual overlay of the tumour regions with standard animal holders used for MSOT and MRI. Finally, we demonstrate the application of the co-registration framework for comparison of perfusion-based data recorded using MSOT and MRI.

## 2. Methods

### 2.1. Animal Experiments

All animal procedures were conducted in accordance with project (70-8214) and personal license (IDCC385D3) issued under the United Kingdom Animals (Scientific Procedures) Act, 1986 and were approved locally under compliance form number CFSB0671. Subcutaneous tumours were established in male BALB/c nude mice (Charles River, 7-10 weeks old, 17-22g) by inoculation of cells from one of three different cancer cell lines in both flanks (1.5×10^6^ LNCaP prostate adenocarcinoma cells, n=3 mice; 1.5×10^6^ PC3 prostate adenocarcinoma cells, n=3 mice; 1×10^6^ mouse K8484 pancreatic adenocarcinoma cells, n=3 mice) in 100*µ*L phosphate buffered saline (PBS). Using three different cell lines allowed us to investigate the co-registration procedure across a range of morphological and functional characteristics.

### 2.2. Multispectral Optoacoustic Tomography (MSOT)

An MSOT inVision 256-TF commercial small animal imaging system (iThera Medical GmbH) was used. Briefly, a tunable optical parametric oscillator (OPO) pumped by an Nd:YAG laser provides excitation pulses with a duration of 9 ns at wavelengths from 660 nm to 1200 nm at a repetition rate of 10 Hz with a wavelength tuning speed of 10 ms and a peak pulse energy of 90 mJ at 720 nm. Ten arms of a fibre bundle provide uniform illumination of a ring-shaped light strip of approximately 8 mm width. For ultrasound detection, 256 toroidally focused ultrasound transducers with a centre frequency of 5 MHz (60% bandwidth) are organized in a concave array of 270 degree angular coverage and a radius of curvature of 4cm.

Mice were prepared according to our standard operating procedure [15]. Each mouse was anaesthetised using <3% isoflurane and moved into a custom animal holder (iThera Medical GmbH), wrapped in a thin polyethylene membrane, with ultrasound gel (Aquasonic Clear, Parker Labs) used to couple the skin to the membrane. The holder was then placed within the MSOT system and immersed in degassed water maintained at 36 °C. The mouse was allowed to stabilise for 15 minutes within the system prior to initialisation of the scan and its respiratory rate was then maintained in the range 70-80 bpm with ∼ 1.8% isoflurane concentration for the entire scan. The imaging slice was chosen to show largest cross-sectional area of the tumours on one or both flanks where possible. Images were acquired in the single slice using 10 wavelengths between 700 nm and 880 nm and averaging of signals from 6 pulses per wavelength; a single slice acquisition was 5.5s in duration. For Oxygen Enhanced Optoacoustic Tomography, 70 such images were acquired continuously, with the breathing gas switched from medical air (21% Oxygen) to pure oxygen (100% Oxygen) after 30 scans, for the purpose of quantification of the response in blood oxygen saturation to such defined oxygen challenge.

### 2.3. Magnetic Resonance Imaging (MRI)

A 9.4 T Agilent MRI system (Agilent, Santa Clara, USA) running VnmrJ 3.1, using an Agilent quadrature transmit/receive millipede volume coil of 38 mm inner diameter was used. The same anaesthesia protocol as for optoacoustic imaging experiments was maintained. A physiological monitoring system was used for observing mouse status and for sequence triggering (SAII, Stony Brook, NY, USA). The core temperature of the mouse was monitored using a rectal probe, and stabilized to 37 °C using an air heating system. Axial multislice T2-weighted images were acquired covering the entire tumour using a respiratory-gated fast spin-echo sequence (field of view 40mm, slice thickness/gap 0.95/0.05mm, TR 2000ms, echo spacing 9ms, echo train length 8, effective TE 36ms, 2 averages, 3-4 slices acquired per gate) with chemical-shift-selective fat suppression.

Dynamic contrast enhanced (DCE)-MRI data were acquired using a spoiled gradient echo sequence (field of view 40mm, 2mm slice, 128×128 points, TR 20ms, TE 1.62ms, 2 averages). 10 images were acquired during the 1 minute prior to administration of contrast agent (Gadavist, Bayer, 200*µ*mol/kg) to provide a baseline reference and 120 images were acquired in the 11 minutes after injection.

### 2.4. Hardware Co-registration

To facilitate co-registration of MSOT and MRI data, a new small animal holder was developed to reproduce the spatial positioning and body deformation of the MSOT (Figure 1a) during the MRI acquisition as accurately as possible. This was achieved using a silicone bed (Figure 2a), fabricated based on photogrammetry of a mouse suspended in PE film, performed with the software 3DF Zephyr v3.5 (3DFLOW, Italy). The deformation in the resulting 3D model was transferred to an isosurface extraction from the Digimouse atlas and subsequently converted into a 3D model (Figure 2b). The resulting model was converted into a negative mold in STL file format, then printed with Polylactic acid (PLA) using an Anet A6 3D printer (Anet, China), instructed with the slicer software Ultimaker Cura 2.6. The 3D printed mold was inserted into a conventional MRI bed and the resulting cavity was filled with silicone (Polycraft T15 Translucent Silicone, MB Fibreglass) before being cured for 24 hours. Subsequently, the negative mold was removed, the silicone bed taken out and the excess silicone trimmed.

**Figure 1:**
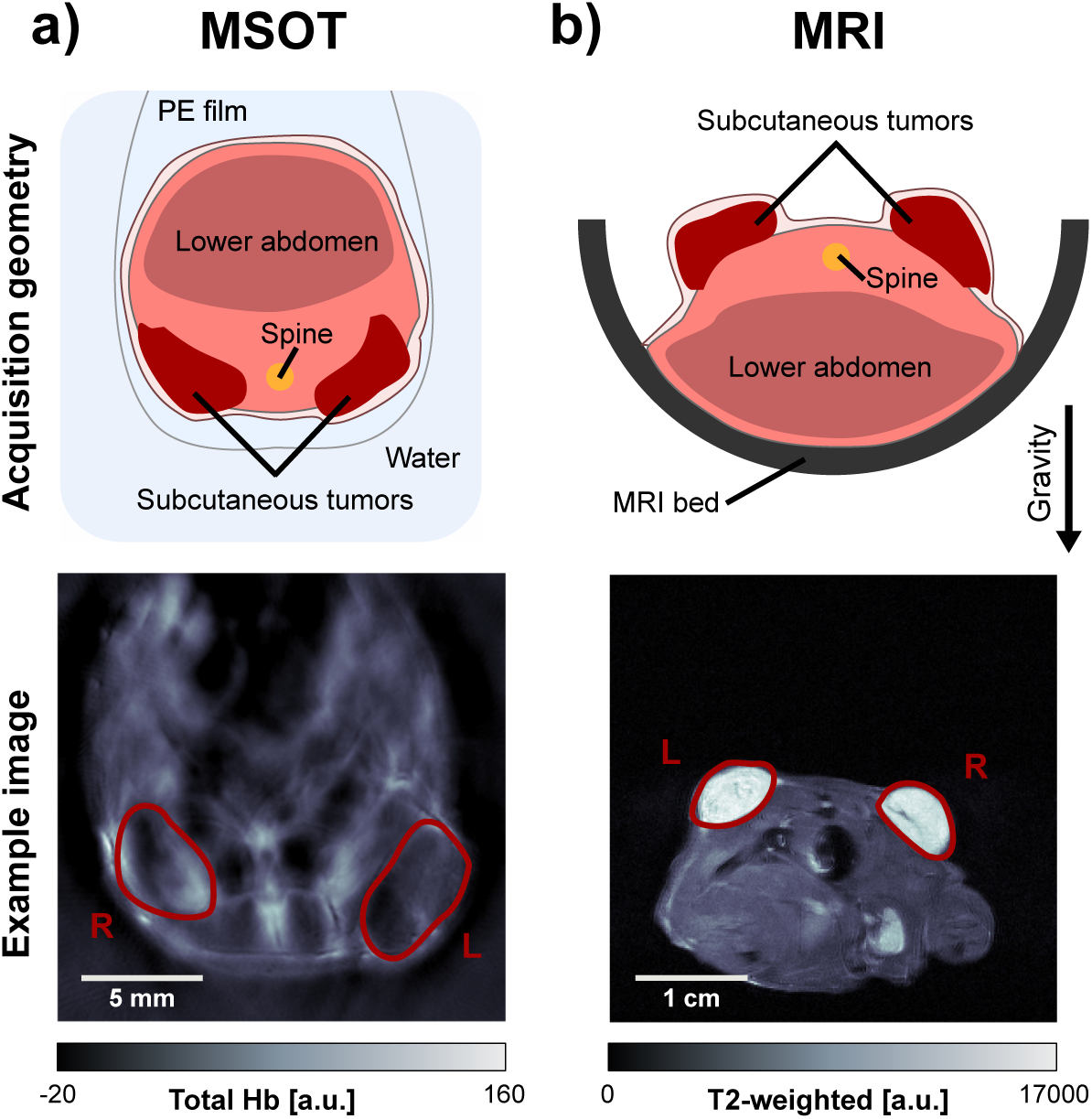
Conventional MSOT and MRI holder geometries. (a) Animal holder geometry and example image showing the total haemoglobin signal after spectral unmixing, acquired using MSOT. (b) Conventional animal holder geometry and example fast spin-echo image from MRI.

**Figure 2:**
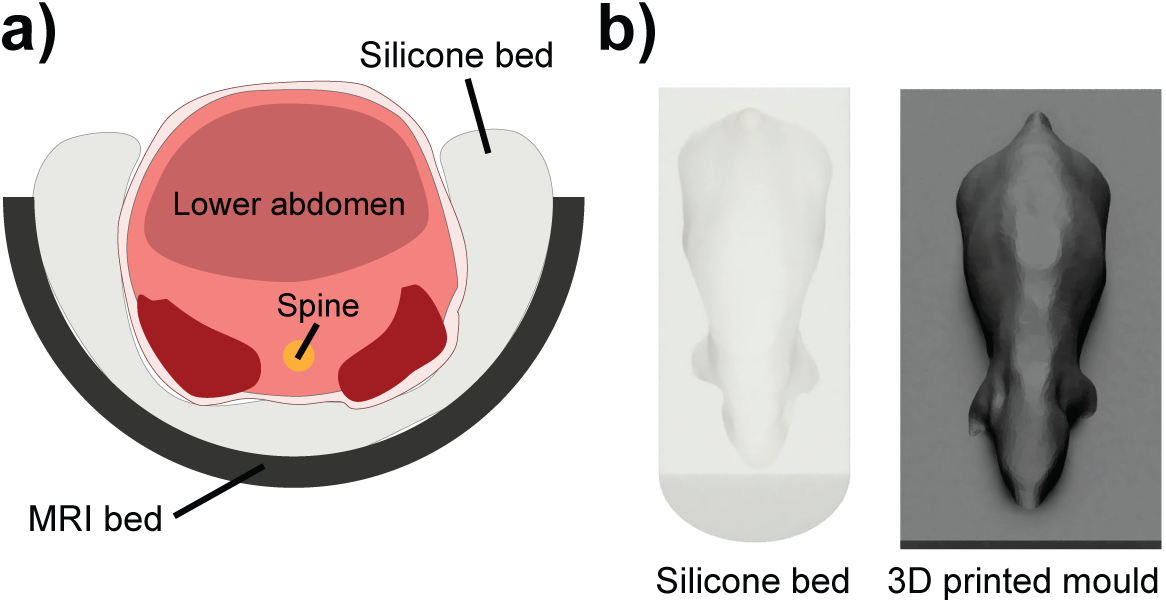
Novel MRI holder geometry. (a) Design of the silicone holder to achieve a comparable anatomical cross-section within MRI and MSOT. (b) Rendering of the silicone bed and the corresponding 3D printed mold.

After imaging in the MSOT, mice were maintained under anaesthesia and transferred for MRI. A subset of n=4 mice (3 LNCaP tumour bearing and 1 PC3 tumour bearing) underwent MRI placed in the prone position in a half-pipe plastic holder, with the tumours on the back facing upwards using the conventional MRI holder geometry (Figure 1 b).

The remaining 5 animals (2 PC3 tumour bearing and 3 K8484 tumour bearing) were scanned in the custom silicone holder (Figure 1 c,d). Transfer into the silicone MRI bed was made in a smooth motion while maintaining the supine orientation of the mouse to preserve the positioning. The silicone bed showed a large, broad nuclear magnetic resonance excitation at 7ppm upfield of water, which was clearly visible in fast spin-echo images. Image registration was greatly simplified by suppressing this signal, as the silicone is not present in the MSOT image data. Therefore, for imaging sessions employing the silicone bed, a modified chemical-shift selective fat suppression sequence was employed, using a sinc pulse of bandwidth 3kHz centred at 2kHz from the water peak, between the fat and silicone resonances, to suppress both fat and silicone.

### 2.5. Software Co-registration

The main objective of any general co-registration software is to merge the coordinate system of the moving image *I*_*M*_ with the fixed (or reference) image *I*_*F*_. The transformation matrix T is used to warp the moving image in order to minimise the error metric with the fixed image. This process is iterated until a certain convergence criterion is reached.

A landmark-based co-registration approach [16] based on non-reflective similarity with the addition of optional reflection was used to register the tumour areas between modalities, utilising a set of prominent anatomical features including the tumour edges and spine location as landmarks. The positions of these features, identified manually, were denoted in the MR image as *I*_*M*_, and the matching positions in the MSOT image as *I*_*F*_. These vectors were then used to calculate the transformation matrix and transform both modalities into the same coordinate space, minimising the euclidean distance between the landmarks.

The registration procedure was implemented in two steps. The first step ensured proper alignment between the body contours in MSOT and MRI, while the second step provided further alignment of the tumours. Landmarks for the first step were the spine and characteristic anatomical features visible in MRI and MSOT, such as contact points between tumors and body (Figure 3). Second step landmarks were defined by points along the outline of the tumour: up to two points on an axis between the tumour and mouse body; and up to two points on the perpendicular axis (Figure 5). The corresponding similarity-based transformation matrix was calculated using the MATLAB function *fitgeotrans*.

**Figure 3:**
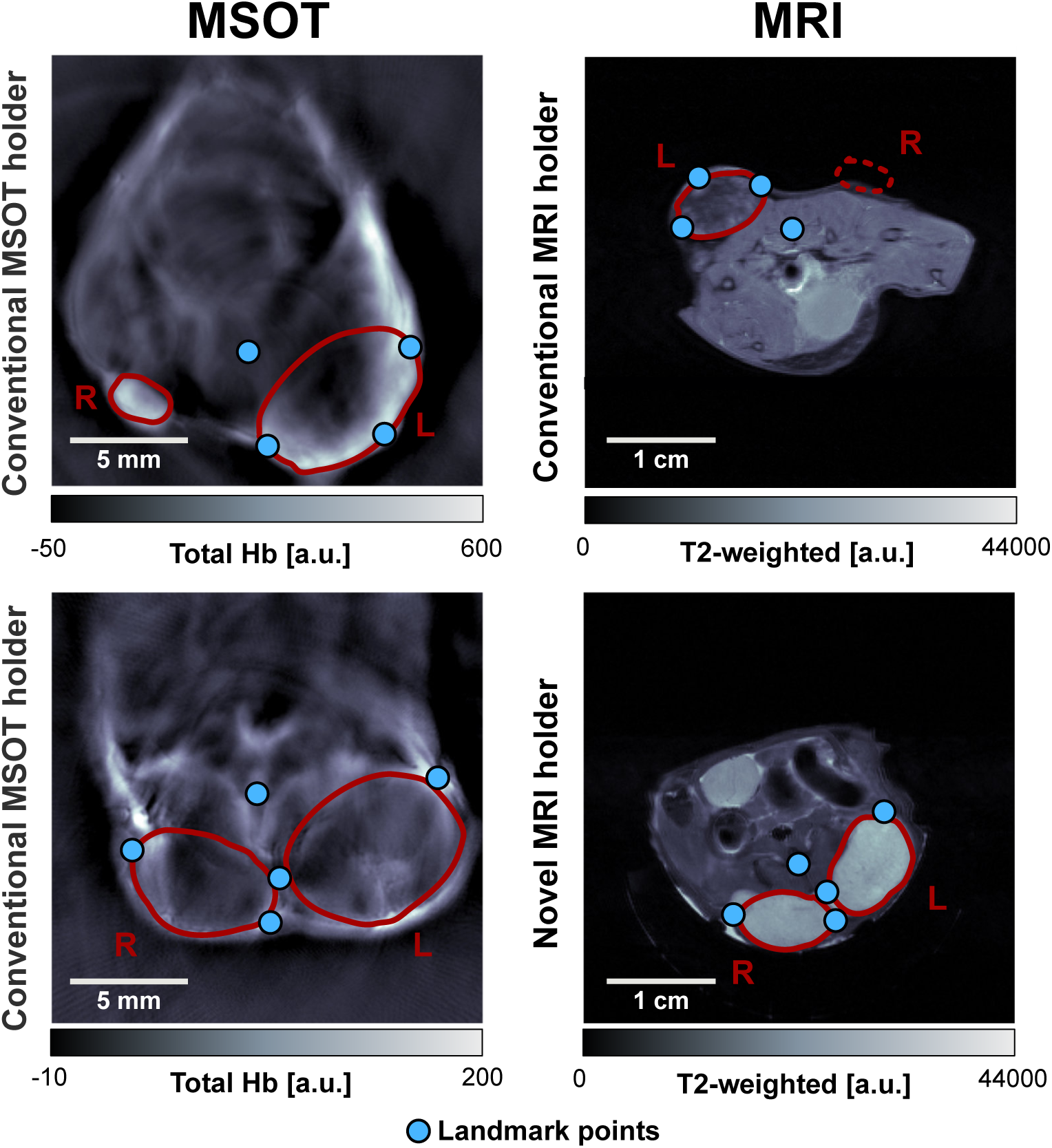
Qualitative comparison of tomographic image data from OT and MRI modalities using the conventional and novel MRI holders. *Top*: MSOT/MRI image pair with conventional holder geometry as in Figure 1. Tumour R is not visible in the corresponding MR image. *Bottom*: MSOT/MRI image pair with novel holder geometry as in Figure 2.

### 2.6. Image and Statistical Analysis

All image analysis was performed in MATLAB 2016a (Mathworks) using the Image Processing Toolbox, the Computer Vision Toolbox and custom scripts unless otherwise stated. All image data and custom analysis codes will be made openly available at doi: 10.17863/CAM.39741.

Image reconstruction was performed using an acoustic backprojection algorithm (iThera Medical GmbH) with an electrical impulse response correction, to account for the frequency dependent sensitivity profile of the transducers. Images were reconstructed with a pixel size of 100 µm × 100 µm which is approximately equal to half of the in-plane resolution of the InVision 256-TF. Pseudoinverse matrix inversion (pinv function in MATLAB 2016a) was applied to the measured optoacoustic spectrum in each pixel to calculate the relative oxy-[*HbO*_2_] and deoxy-haemoglobin [*Hb*] signal. The presented images illustrate the total haemoglobin signal [*HbO*_2_ + *Hb*] unless otherwise stated. Apparent blood oxygen saturation 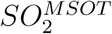 was calculated as a ratio of oxy-to total haemoglobin [4].

All MR images were flipped horizontally prior to image registration. The position of the slice analysed was chosen by the operator to best match the location of the imaging slice in OT, acquired directly before the MRI.

The analysis of registration accuracy of body and tumour contours was performed by calculating the Dice Similarity Coefficient (DSC). This coefficient is defined as:

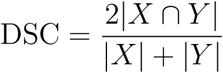

which allows quantification of the overlap between two binary masks, X and Y (i.e. original MRI body/tumour mask and the mask obtained from the MSOT image after co-registration). The higher the DSC, the better the overlap between the two binary masks and therefore, the more accurate the image registration result.

The results were compared on a per-tumour basis. Differences in DSCs between conventional and novel holder geometries as well as before and after landmark-based registrations (for body and tumor) were statistically tested with two-tailed paired t-tests (in the case of equal variances between sets of samples) and two-tailed unpaired t-tests (in the case of unequal variances between sets of samples). Data are reported as median ± standard deviation, unless otherwise stated.

DCE-MRI signal Area under the Curve (AuC) 1 minute after contrast administration was compared to the change in blood oxygen saturation 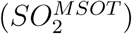 as measured by Oxygen Enhanced Optoacoustic Tomography in response to an oxygen challenge [4]. The median DCE AuC values in the regions showing positive 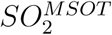 response and the rest of the tumour area were compared, with the pixels classified as positively responding when the difference between the average 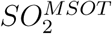 in the first 20 frames (under air breathing) and the the last 20 frames (oxygen breathing), exceeded twice the standard deviation of the 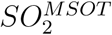 in the first 20 frames.

## 3. Results

### 3.1. Novel holder geometry improves visual anatomical similarity between MSOT and MRI

Visual inspection of MRI and MSOT images acquired with the conventional and novel MRI holder geometries yielded distinct differences in body shape and anatomical appearance (Figure 3). Overall body shape and relative tumour location were not easily comparable for tumours imaged with the conventional protocol, while the novel protocol showed a high degree of similarity in body contour and tumour locations. A quantitative comparison of the contours of the mouse bodies in MSOT and MRI images using Dice similarity coefficients (DSCs) showed significant improvement (p=0.03, unpaired t-test) with the novel MRI holder, resulting in a higher DSC (0.63 ± 0.05 vs. 0.52 ± 0.07, novel vs. conventional). The higher DSC indicates that the novel MRI holder more accurately represents the body deformation observed in the MSOT.

### 3.2. Landmark-based contour registration improves animal body alignment

Landmark-based contour registration was then applied to MSOT and MR images acquired using both the conventional and novel MRI holders (illustrated in Figure 4a). The difference between the conventional and novel MRI holders was more significant (p=0.002, unpaired t-test) following landmark-based registration with further improved DSCs (0.92 ± 0.02 vs. 0.83 ± 0.03, novel vs. conventional). In total (Figure 4b), the contour registration procedure improved DSCs for body contour overlay significantly, both for the conventional (ΔDSC = 0.31, p=0.004, paired t-test) and novel (ΔDSC = 0.29, p=7.2 ×10^−5^, paired t-test) holder.

**Figure 4:**
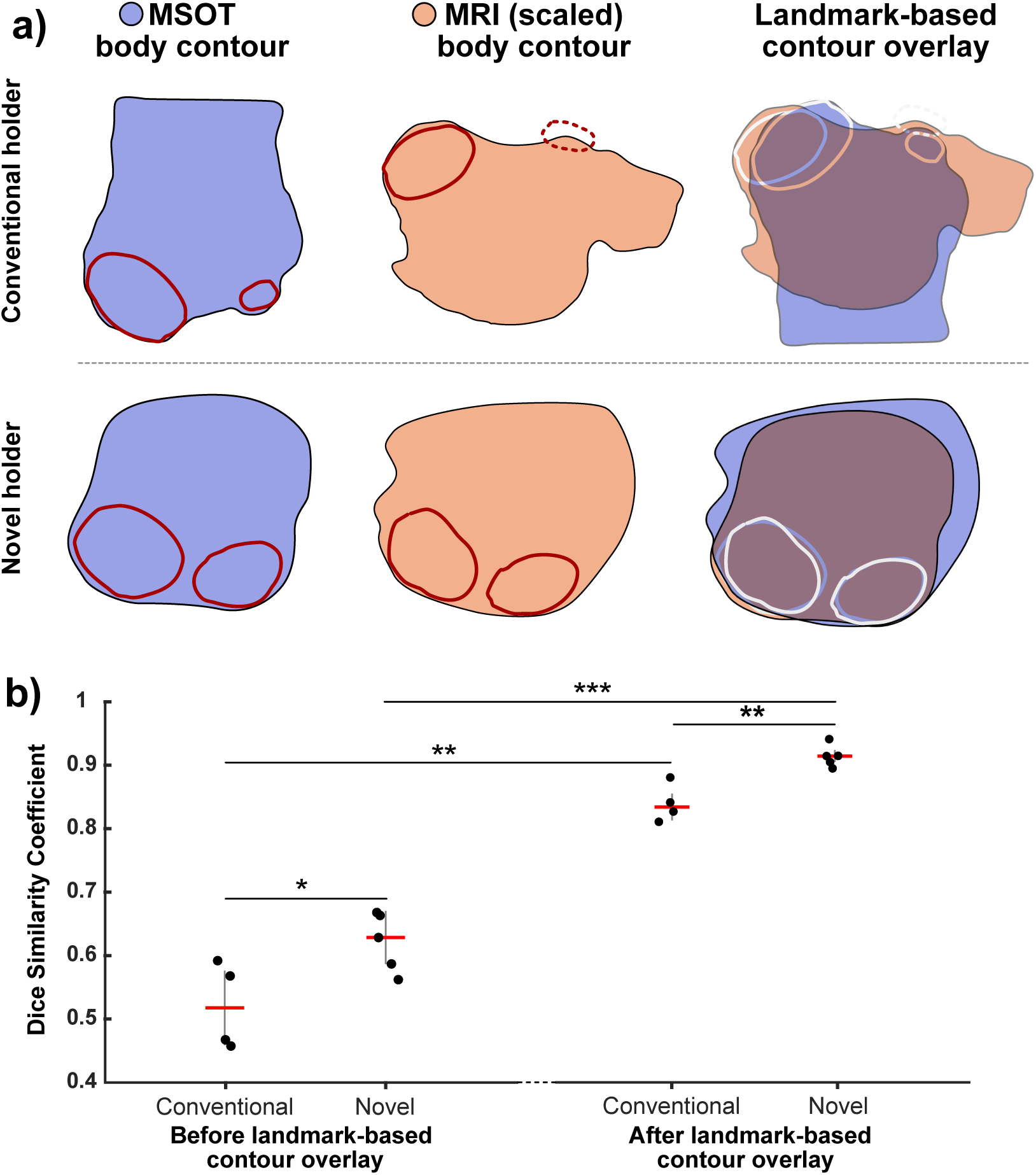
Comparison of body contour overlays of MRI/MSOT image pairs for conventional and novel holder geometries. (a) Body contour pairs (derived from binarized body outlines) for MSOT and MRI. Use of the novel protocol improves the agreement substantially (overlaid MSOT and MRI contours shown in blue and orange respectively).(b) Quantitative comparison of Dice similarity coefficient (n=4 for conventional holder and n=5 for novel MRI holder). * p<0.05, ** p<0.01, *** p<0.001 by unpaired two-tailed t-test (unequal variances) and paired two-tailed t-test (equal variances).

### 3.3. Landmark-based tumour contour optimisation further improves local anatomical similarity

Following the body contour registration, each tumour was individually co-registered as an additional optimisation step. The tumour contours showed a qualitatively higher agreement after this additional landmark-based optimisation. The gain in registration accuracy was estimated to be between 5 and 15 pixels (375 µm - 1125 µm), based on the distances between the co-registered tumour outlines (Figure 5a). Quantitative assessment resulted in a significant improvement in tumour mask overlay DSCs (p=0.005, paired t-test) after landmark-based transformation of tumour masks (pre-transform: 0.85 ± 0.06 vs. post-transform: 0.92 ± 0.04, Figure 5b).

**Figure 5:**
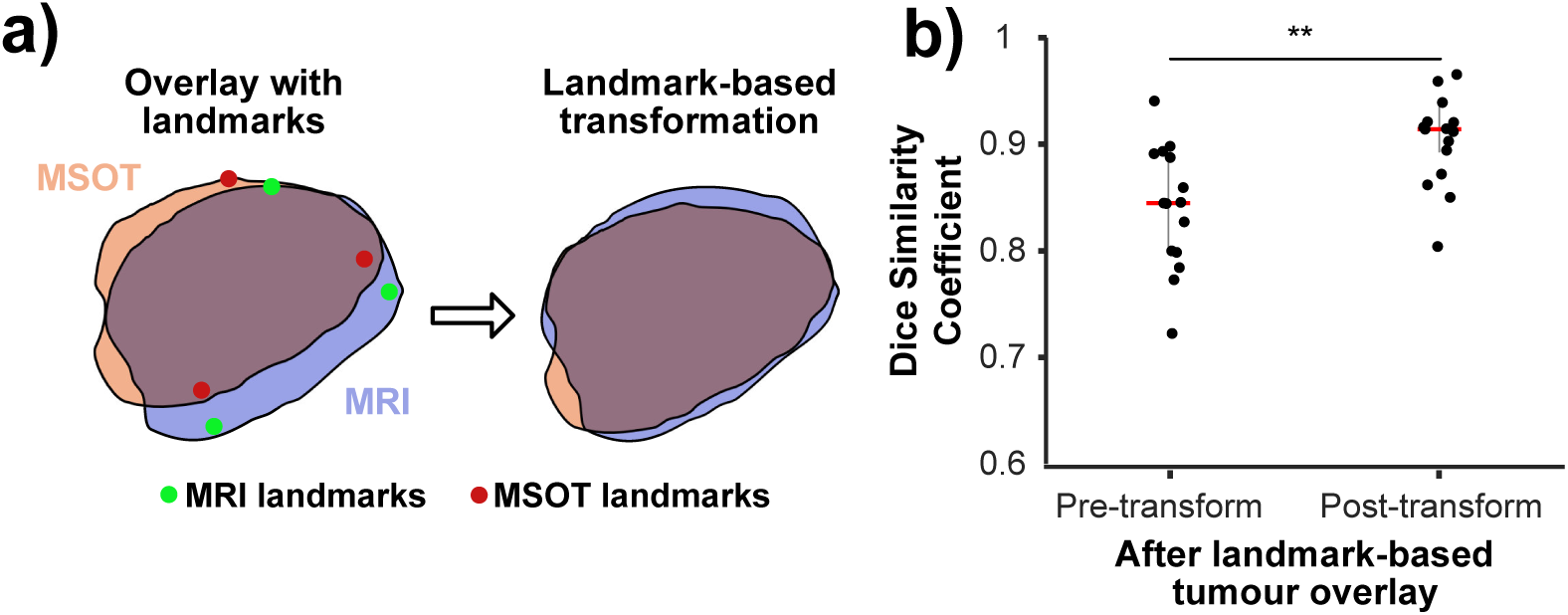
Overlays of tumour contours from MRI/MSOT image pairs before and after landmark-based optimization. (a) Comparison of a tumour outline before and after landmark-based tumour registration. (b) Quantification of the improvement in Dice similarity coefficient (n=15 tumours, combined data for conventional and novel MRI holder geometry). ** p<0.01 by paired two-tailed t-test (equal variances).

### 3.4. Application of the co-registration framework for comparison of data acquired using MSOT and MRI

Comparison of the anatomical similarity of the imaging data from the two modalities subjected to our co-registration framework was made in three K8484 tumour bearing mice. K8484 tumours were used for this purpose as they contain heterogeneous structural features visible in both MSOT and MRI. Upon visual inspection of images from three mice bearing this tumour type, it can be seen that the body shapes and tumour locations images demonstrate high anatomical similarity (Figure 6). Considering the feature locations (defined as distinct features in MRI and MSOT images belonging to the same structure, denoted by red annotations in Figure 6), we established that the relative distance between the centres of the features between modalities showed a close agreement (2,4, and 12 pixels, or 150, 300 and 900 *µ*m respectively for the three mice shown). In Figure 6, the red rectangle indicates the extent of the observed features in MRI/MSOT image pairs, whereas the red asterisk highlights the most distinct point within the feature. This point was subsequently used for determining the relative distance between modalities.

**Figure 6:**
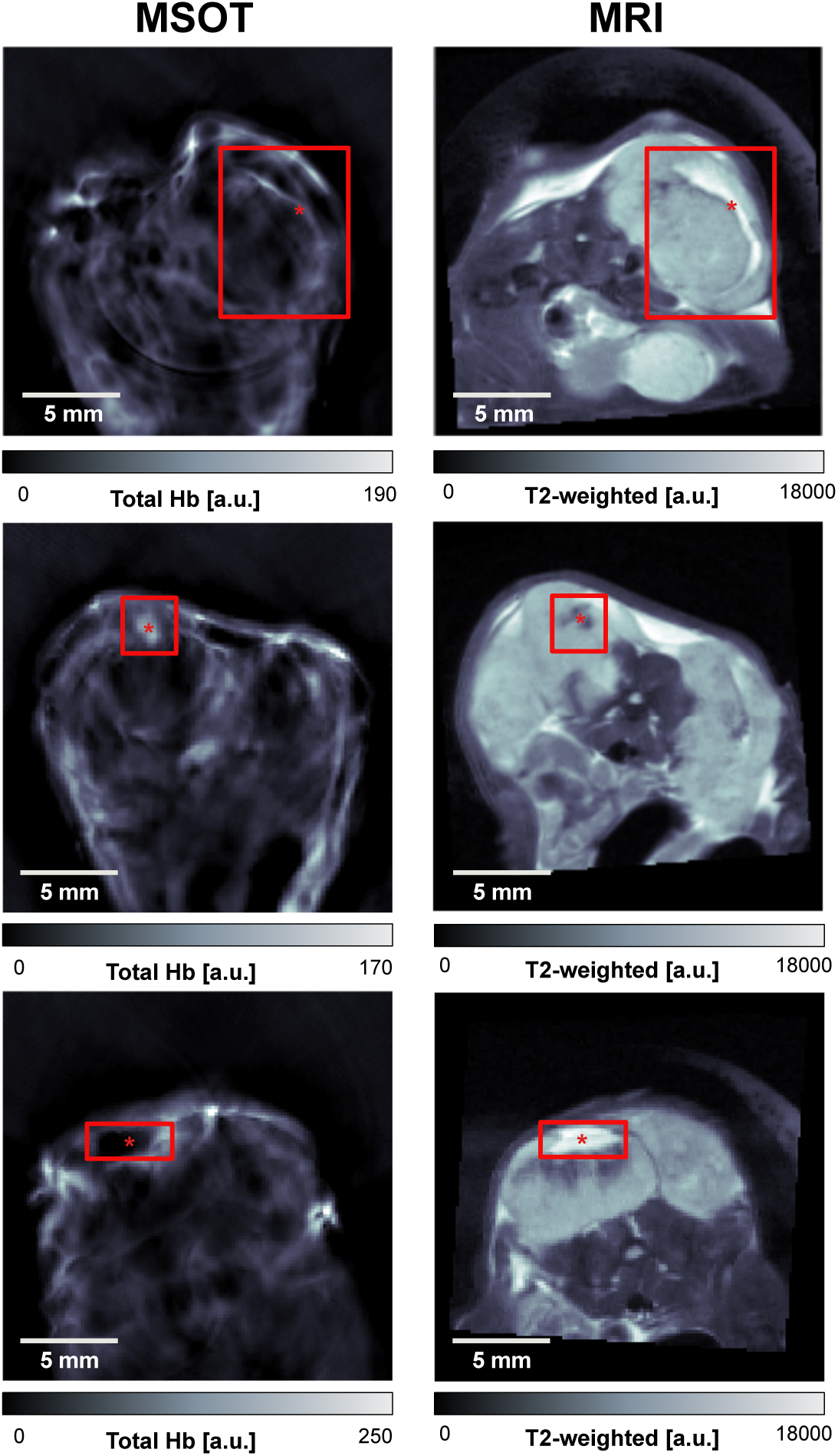
Anatomical features in three K8484 tumours after landmark-based body and tumour contour registration. All tumours contained distinct intratumoural features (highlighted in red). The distances between the estimated feature centroids (marked with red asterisks) were measured to be very small after registration, with a range from 2 to 12 pixels (150 µm to 900 µm).

A comparison of functional imaging data was then made in these K8484 tumours based on imaging data recorded using DCE-MRI and Oxygen Enhanced OT (OE-OT) protocols, which have been previously shown to relate to tumour perfusion and vascular function [6]. Visual comparison of DCE-MRI and OE-OT images (Figure 7a) shows a similar distribution of perfused pixels in both modalities, with a greater number in the rim compared to the core of the tumour, as is commonly reported in subcutaneous xenografts. Quantitative comparison of DCE-MRI enhancement in regions of OE-OT response (Figure 7b) shows a markedly stronger DCE-MRI enhancement in the areas showing positive response in the OE-OT, suggesting a functional relationship between these imaging biomarkers.

**Figure 7:**
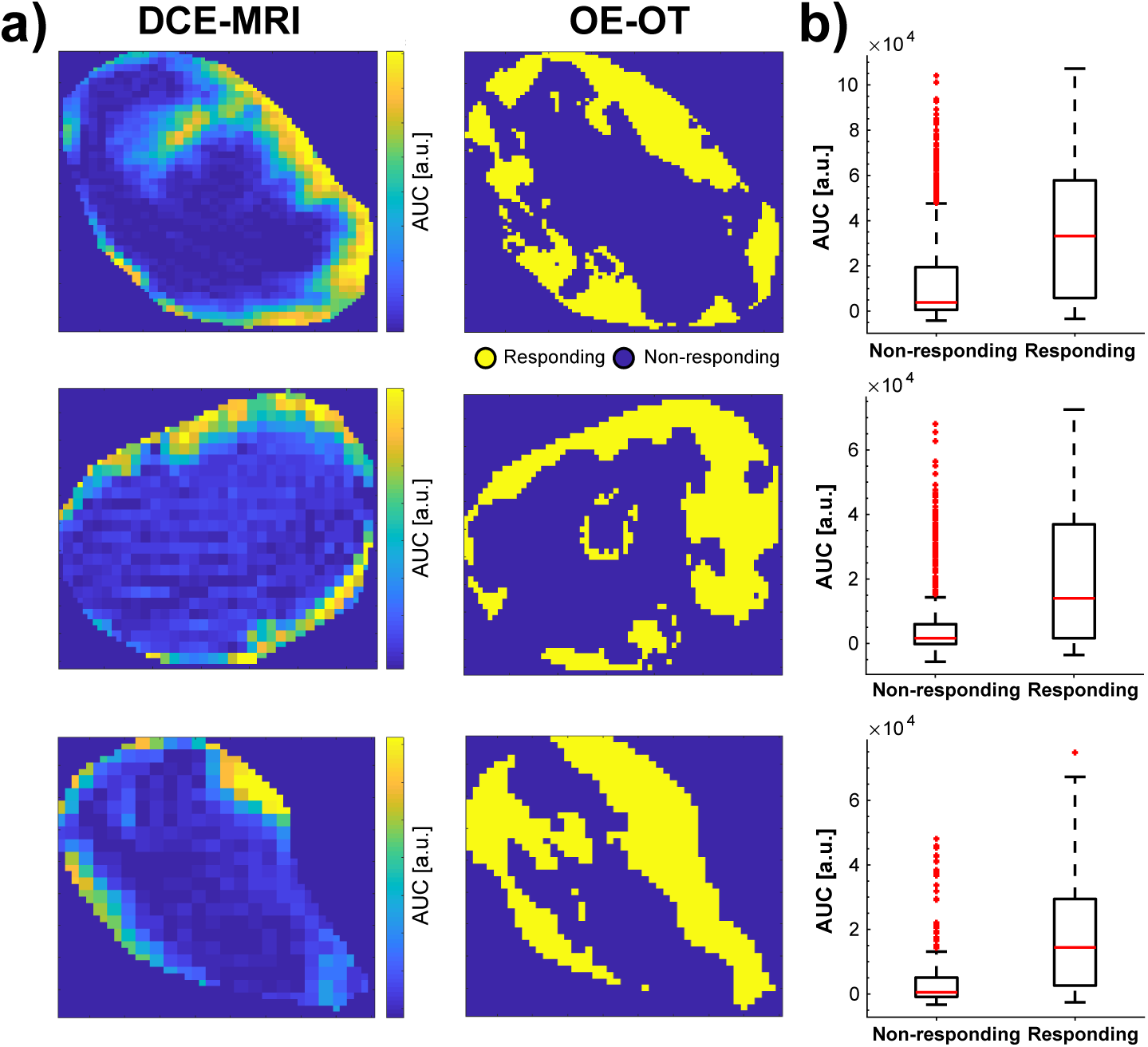
Measurements of vascular function show strong co-localisation in MRI and MSOT. (a) Maps of DCE-MRI area under the curve (AUC) enhancement 1 minute after contrast injection (left) are in close spatial agreement with the maps of positive response to oxygen challenge in Oxygen-Enhanced Optoacoustic Tomography (right). (b) The DCE-MRI AUC is clearly higher in the areas positively responding in OE-OT.

## 4. Discussion

Co-registration of images between modalities enables the combination of complementary information provided by different imaging methods. Due to deformation of the animal or patient between scans, correct alignment of images can pose a significant challenge and require both hardware and software-based optimisation approaches. In this work, we describe an integrated hardware and software framework for co-registration of small animal MSOT and MR imaging data. Without co-registration, these modalities produce very different images of the sample, due to different animal positioning and stress distribution.

On the hardware side, a novel silicone MRI animal holder was developed, which was designed to mimic the external stresses acting on the mouse body in the MSOT. Introducing the new holder alone already significantly improved the similarity in the shape of the entire mouse body contour as well as the individual tumour contour, contributing to more accurate co-registration. Importantly, the use of the holder did not increase animal preparation time or cause any side effects for animal welfare during imaging. Fabrication of the holder is simple and inexpensive, as soft two-component silicone is poured over a 3D printed mouse mold. The protocol offers a simple solution to improve MSOT/MR image co-registration.

A software tool for landmark-based image co-registration was then established to further improve the co-registration and enable per-pixel analysis of the combined multi-modal images. The transformation matrix for the MSOT images was calculated to maximise similarity between body and tumour outlines in both modalities as well as to minimise the distances between anatomical landmarks. The result of applying this software tool was a co-localisation error in the order of 100 microns, comparable to the typical resolution of both modalities. This framework also enabled per-pixel combination and comparison of the insight offered by MSOT and MRI in functional imaging. The relationship between tumour perfusion, provided by early DCE-MRI enhancement [9], and vascular function, given by the MSOT response to oxygen challenge [6], served as a proof of concept for further MSOT/MRI comparison.

Despite the clear improvements in image co-registration achieved, there remain some limitations to our study. Firstly, the described two-step hardware and software framework is designed to aid with 2D co-registration, which assumes already the correct, manual choice of matching imaging slice between the modalities. The use of the silicone holder can help in this task to some extent, as the similar cross-sectional shape of the tumour in the MRI can help match it qualitatively to the geometry in the MSOT. Slice misalignment will introduce additional error in the co-registration procedure.

A second limitation arises in the design of the silicone bed, which aimed to mimic the effects of the polyethylene film holder used in the MSOT, as well as the stresses due to water submersion during MSOT imaging. In order to support the weight of the animal, the silicone had to be stiffer than optimal, causing some discrepancy in MRI/MSOT mouse positioning. Further optimisation using silicones of different elastic properties could better match the distribution of forces and should be investigated in future experiments.

Finally, the silicone bed was created for a specific mouse size based on the typical usage in our experiments. If needed, additional silicone beds could be created to account for different mouse sizes, across strain and age for instance, and taking the individual tumour position into consideration. The optimal approach would utilize 3D modeling to create mouse-specific holders and require standardisation of the modelling, printing and casting workflow.

### 4.1. Conclusion

We have demonstrated the feasibility of a hardware- and software-based image registration framework for MRI and MSOT images. We use a novel silicone MRI holder, as well as a software tool to perform landmark-based co-registration of the images. Both steps led to a significant improvement in the registration of the tumour outlines and internal structure between the modalities. This simple, inexpensive approach can be readily implemented for multi-modal MSOT/MRI studies of small animals, which will help to provide valuable insight into relative performance of these two modalities in revealing vascular architecture and function in cancer.

## 5. Acknowledgements

This work was supported by Cancer Research UK (C47594/A16267, C14303/A17197) and the EPSRC-CRUK Cancer Imaging Centre in Cambridge and Manchester (C197/A16465 and C8742/A18097). We would like to thank the CRUK CI Core Facilities for their support of this work, in particular the Imaging Core, Biological Resource Unit, Histopathology, and Biorepository. We also want to thank Mireia Crispin-Ortuzar and Joanna Brunker for helpful comments on the draft article.

